# Low-dose IFN-γ induces tumor cell stemness in tumor microenvironment of non-small cell lung cancer

**DOI:** 10.1101/517003

**Authors:** Mengjia Song, Yu Ping, Kai Zhang, Li Yang, Feng Li, Shaoyan Cheng, Dongli Yue, Nomathamsanqa Resegofetse Maimela, Jiao Qu, Shasha Liu, Ting Sun, Zihai Li, Jianchuan Xia, Bin Zhang, Liping Wang, Yi Zhang

## Abstract

Interferon-γ (IFN-γ) is conventionally recognized as an inflammatory cytokine that play a central role in antitumor immunity. Clinically, although has been used clinically to treat a variety of malignancies, low-level IFN-γ in the tumor microenvironment (TME) increases the risk of tumor metastasis during immunotherapy. Accumulating evidence has suggested that IFN-γ can induce cancer progression. The mechanisms underlying the controversial role of IFN-γ regulating tumor development remain unclear. Herein, we firstly revealed a dose-dependent effect of IFN-γ in inducing tumor stemness to accelerate cancer progression in patients with a variety of cancer types. Mechanically, low-level IFN-γ endowed cancer stem-like properties via the intercellular adhesion molecule-1 (ICAM1)-PI3K-Akt-Notch1 axis, whereas high-level IFN-γ activated the JAK1-STAT1-caspase pathway to induce apoptosis in non-small cell lung cancer (NSCLC). Inhibition of ICAM1 abrogated the stem-like properties of NSCLC cells induced by the low dose of IFN-γ both *in vitro* and *in vivo*. Our study first defines the role of low-level IFN-γ in conferring tumor stemness and clearly elucidate the distinct signaling pathways activated by IFN-γ in a dose-dependent manner, providing new insights into cancer treatment, particularly patients with low-level IFN-γ expression in the TME.

## Introduction

Interferon (IFN)-γ is widely considered to be a crucial antitumor cytokine that is mainly produced by activated T cells, natural killer (NK) and NKT cells(Sadanaga et al, 1999; Tannenbaum & Hamilton, 2000). After binding to the heterodimeric IFNGR1/IFNGR2 receptor complex, IFN-γ initiates the activation cascade of downstream signaling events, particularly classical JAK-STAT signaling, and the transcription of multiple IFN-γ-inducible genes, both of which induce cell cycle arrest and apoptosis in tumor cells(Platanias, 2005). However, IFN-γ signaling has paradoxically been reported to promote carcinogenesis and metastasis(Muller-Hermelink et al, 2008; Murohashi & Hoang, 1991) through inducing inflammatory responses(Xiao et al, 2009), immunosuppression or other unknown mechanisms(Katz et al, 2008; Ostrand-Rosenberg & Sinha, 2009). Sustained low-level IFN-γ boosted the development of several types of tumor in mouse models(He et al, 2005; Kelly et al, 1991). Clinically, although IFN-γ has been used in many anti-cancer clinical trials, low-level IFN-γ generated at the tumor site has been shown to increase the risk of tumor metastasis during immunotherapy(Gong et al, 2008). Unfortunately, the molecular mechanisms underlying IFN-γ mediating cancer progression were not fully characterized in these previous studies. Notably, recent studies have described a close relationship between IFN-γ and cancer stem cells (CSCs). IFN-γ are capable of inducing CSCs dormancy(Liu et al, 2017) and metastatic CSCs generation(Chen et al, 2011). These studies imply that IFN-γ may involve in tumor progression through conferring tumor stemness, but the exact mechanism was not fully understood.

CSCs have been reported to play key roles in tumor initiation, metastasis, recurrence and multidrug resistance in various cancer types(Batlle & Clevers, 2017). CSCs are a small subset of self-renewed, pluripotent, immune-privileged, high-tumorigenic and long-living malignant cells. Persistent activation of highly conserved signaling pathways, such as the Notch, Hedgehog, and/or Wnt pathways, partially determines the stem-like properties and tumorigenicity of CSCs(Takebe et al, 2015). In addition, epithelial-to-mesenchymal transition (EMT), an essential developmental process that is often activated during cancer invasion and metastasis(Mani et al, 2008), was also verified as a characteristic of CSCs. Recently, a new concept of “CSC immunology” has been established indicating that the features of CSCs are dependent on a specialized immune-niche. Many immune factors produced in tumor niches, such as IL-6, IL-8 and TGF-β, endow tumor cells with stem-like properties to accelerate tumor progression(Plaks et al, 2015; Qiao et al, 2017).

Intercellular adhesion molecule 1 (ICAM1), a transmembrane molecule and member of the immunoglobulin superfamily of proteins expressed by leukocytes, endothelial cells and epithelial cells, is involved in many important processes, including leukocyte endothelial transmigration, cell signaling, cell-cell interaction, cell polarity and tissue stability(Duperray et al, 1997; Kanters et al, 2008; Lawson & Wolf, 2009; Millan et al, 2006). ICAM-1 can be upregulated by lipopolysaccharide and some inflammatory cytokines, such as IFN-γ and TNF-α(Chen et al, 2000; Rothlein et al, 1988). The protumor role of ICAM1 has been identified in various cancers(Huang et al, 2017; Liu et al, 2013; Min et al, 2017; Tsai et al, 2015), which has been recently used as a target of engineered chimeric antigen receptor T cell therapy in advanced thyroid tumors(Min et al, 2017) and a marker of CSCs in hepatocellular carcinoma (HCC) and esophageal squamous cell carcinoma (ESCC)(Liu et al, 2013; Tsai et al, 2015). In lung cancer, ICAM1 is highly expressed in CSCs(Levina et al, 2008) and responsible for the cancer initiation and metastasis(Lin et al, 2006). These studies support a potential role of ICAM1 in modulating tumor stemness and progression.

In this study, we firstly found a dose-dependent effect of IFN-γ on tumor stemness in non-small cell lung cancer (NSCLC), ESCC, colorectal cancer (CRC) and HCC patients, indicating that low-level IFN-γ in tumor microenvironment induces cancer stem properties. Then, we elucidated the molecular mechanisms underlying this effect in NSCLC cells and identified that low-level IFN-γ facilitated the stem-like properties through the ICAM1-PI3K-Akt-Notch1 signaling cascade, while high-level IFN-γ induced cell apoptosis through the JAK1-STAT1-caspase pathway. Moreover, the increased stem-like properties of NSCLC cells induced by low-dose IFN-γ were abrogated by inhibition of ICAM1 both *in vitro* and *in vivo*. These findings identify the controversial role of IFN-γ in tumor progression and clearly elucidate the distinct mechanisms underlying IFN-γ influencing tumor stemness in a dose-dependent manner.

## Results

### Low-level IFN-γ expression is closely associated with poor prognosis and increased expression of CSC markers in patients with NSCLC

IFN-γ is known as a representative antitumor cytokine in the tumor microenvironment (TME)(Tannenbaum & Hamilton, 2000). We firstly detected the source of IFN-γ in 5 fresh tumor tissues from patients with NSCLC by flow cytometry. CD3^+^CD8^+^ T cells, CD3^+^CD4^+^ T cells, NK cells and NKT cells showed high production of IFN-γ while CD3^+^CD4^+^Foxp^+^ Treg cells showed low production (Fig. S1A, B). Next, we explored the actual distribution pattern of IFN-γ in 86 NSCLC tissues by immunohistochemistry (IHC) staining. Samples were grouped into ‘low (− or +)’ and ‘high (++ or +++)’ according to the IHC staining score (Fig. 1A). Interestingly, we found that low-level IFN-γ expression accounted for the majority in NSCLC tissues (Fig. 1b). Moreover, low-level IFN-γ expression was strongly correlated with the tumor node metastasis (TNM) stage, brain metastasis, chemoresistance (Table 1) as well as shorter overall survival (OS) and progression free survival (PFS) times (Fig. 1C-E). These results suggest that low-level IFN-γ in the TME may play a role in NSCLC progression.

**Table 1.** Association of IFN-γ expression with clinicopathological features in patients with NSCLC.

| Characteristic | Total | IFN-γ expression (n=86) | | $\chi^2$ | P-value |
| --- | --- | --- | --- | --- | --- |
|  |  | Low<br>n=55(64%) | High<br>n=31(36%) |  |  |
| Age, years |  |  |  |  |  |
| $\geq 55$ | 61 | 40 | 21 | 0.239 | 0.6250 |
| <55 | 25 | 15 | 10 |  |  |
| Gender |  |  |  |  |  |
| Male | 60 | 38 | 22 | 0.03311 | 0.8556 |
| Female | 26 | 17 | 9 |  |  |
| Smoking History |  |  |  |  |  |
| Yes | 45 | 23 | 22 | 6.753 | **0.0094\*\*** |
| No | 41 | 32 | 9 |  |  |
| Differentiation |  |  |  |  |  |
| Well/Moderate | 66 | 41 | 25 | 0.4133 | 0.5203 |
| Poor | 20 | 14 | 6 |  |  |
| Histology |  |  |  |  |  |
| Adenocarcinoma | 47 | 31 | 16 | 0.1805 | 0.6709 |
| Squamous cell carcinoma | 39 | 24 | 15 |  |  |
| TNM stage |  |  |  |  |  |
| I-II | 63 | 36 | 27 | 4.74 | **0.0295\*** |
| III-IV | 23 | 19 | 4 |  |  |
| Recurrence |  |  |  |  |  |
| Yes | 14 | 10 | 4 | 0.4053 | 0.5244 |
| No | 72 | 45 | 27 |  |  |
| Lymph node metastasis |  |  |  |  |  |
| Yes | 29 | 17 | 12 | 0.5398 | 0.4625 |
| No | 57 | 38 | 19 |  |  |
| Bone metastasis |  |  |  |  |  |
| Yes | 13 | 8 | 5 | 0.03875 | 0.8440 |
| No | 73 | 47 | 26 |  |  |
| Brain metastasis |  |  |  |  |  |
| Yes | 7 | 7 | 0 | 4.295 | **0.0382\*** |
| No | 79 | 48 | 31 |  |  |
| EGFR mutation |  |  |  |  |  |
| Yes | 34 | 25 | 9 | 2.237 | 0.1348 |
| No | 52 | 30 | 22 |  |  |
| Chemoresistance |  |  |  |  |  |
| Yes | 20 | 17 | 3 | 5.007 | **0.0252\*** |
| No | 66 | 38 | 28 |  |  |

EGFR, epidermal growth factor receptor.
IFN-γ , interferon gamma.
NSCLC, non-small cell lung cancer.
\* $P < 0.05$ , \*\* $P < 0.01$ .

**Figure 1.**
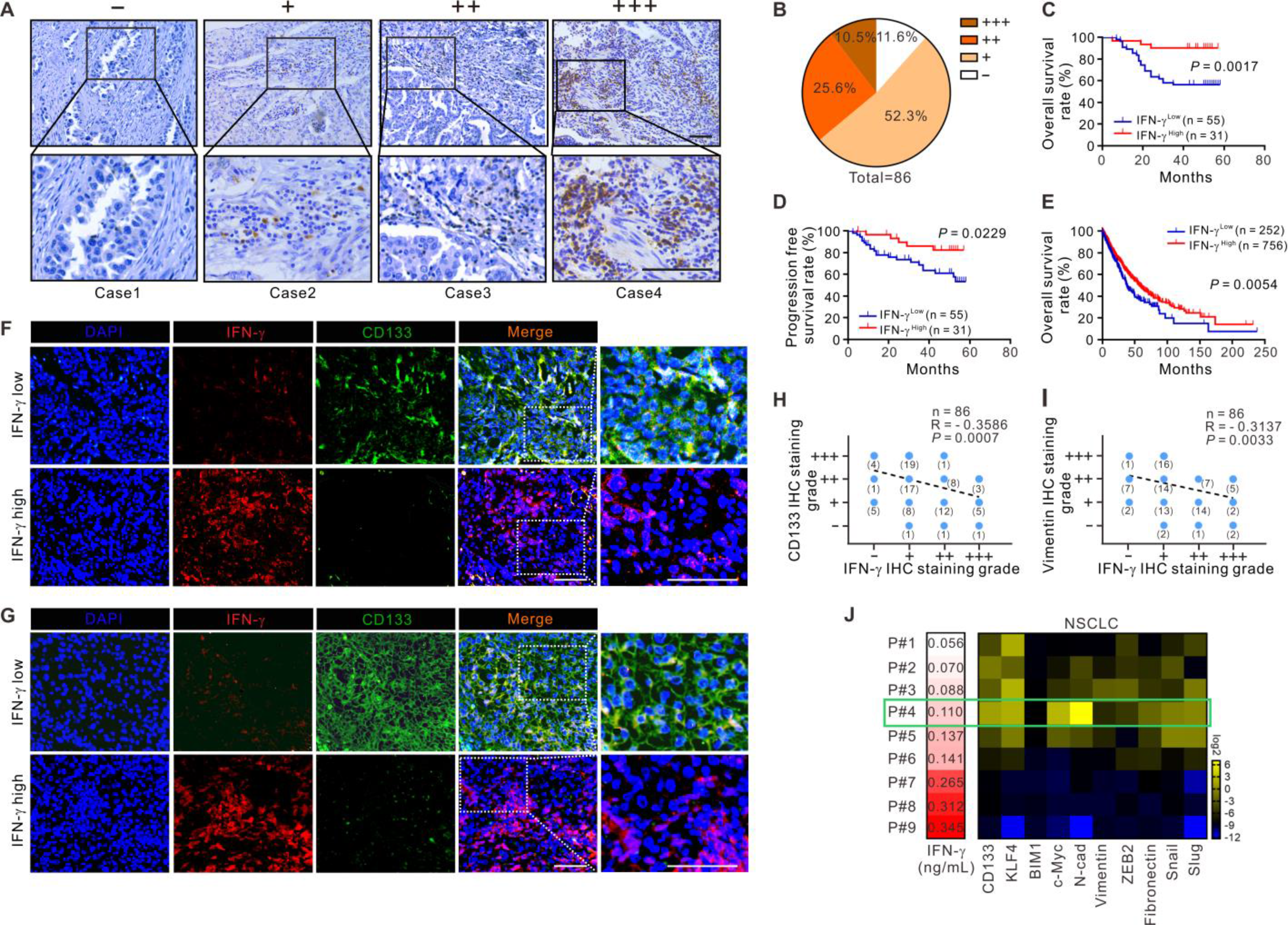
Low-level IFN-γ expression is correlated with tumor stemness in NSCLC patients. **(A)** Representative photomicrographs of IHC staining for IFN-γ in NSCLC tissues. Scale bars, 100 μm. **(B)** Real distribution of IHC results of IFN-γ in 86 NSCLC tissues. **(C, D)** Kaplan-Meier analysis of overall survival and progression free survival in 86 NSCLC patients according to different IFN-γ levels in NSCLC tissues by IHC staining. **(E)** mRNA profiles and follow-up data for 1008 NSCLC patients from TCGA used for analyzing the correlation of IFN-γ mRNA expression and overall survival. **(F, G)** IF staining of the colocalization of IFN-γ (red) with CD133 (green) or Vimentin (green) in NSCLC tissues. Colocalization is shown in yellow. DAPI, blue. Figure panel pairs represent images taken with different zooming options; Scale bars, 100 μm. **(H, I)** Spearman’s correlation analysis was performed to analyze the correlation between IFN-γ and CD133 and Vimentin according to the IHC staining results. **(J)** Heat map showed the expression of IFN-γ and stemness related genes. The concentration of IFN-γ in TIF was detected by ELISA (left). The percentage of CD133 positive cells in tumor cells which was labelled by CD326 positive, was detected by flow cytometry and the expression of other stemness related genes was tested by RT-PCR in NSCLC (right). Patient #4 labelled by green frame showed the highest expression of stemness related genes. Log-rank tests were used in **C-E**. The results are representative of three independent experiments.

CSCs have been reported to participate in tumor initiation, metastasis, recurrence and multidrug resistance(Batlle & Clevers, 2017), serving as a key mechanism of tumor progression. CD133 has been identified as a marker of CSCs in NSCLC(Roy et al, 2017; Zakaria et al, 2017). As CSCs are also characterized by enhanced EMT(Mani et al, 2008), Vimentin, an EMT marker, was used as another CSC marker in our study. By immunofluorescence (IF) staining, we intriguingly found much higher expression of CD133 and Vimentin on tumor cells in IFN-γ-low TME than that in IFN-γ-high TME in NSCLC tissues (Fig. 1F, G). A significantly negative correlation between IFN-γ and CD133 or Vimentin was observed by IHC staining analysis in 86 NSCLC tissues (Fig. 1H, I). In addition, a negative correlation between IFNG and PROM1 was also observed in lung adenocarcinoma collected from TCGA (Fig. S2A). To further accurately quantify the level of IFN-γ promoting tumor stemness, we performed enzyme-linked immunosorbent assay (ELISA) to detect the concentration of IFN-γ in tumor interstitial fluid (TIF) isolated from fresh 9 NSCLC tissues. Accordingly, cancer stem properties were evaluated by the percentage of CD133^+^ stem cells in CD326^+^ tumor cells and the expression of stemness related genes. Intriguingly, we found patients with relative low-level (around 0.137ng/mL) IFN-γ in TIF showed increased cancer stem properties, and patient #4 with 0.110 ng/mL IFN-γ showed the highest percentage CD133^+^ stem cells and highest expression of most stemness related genes. However, patients with high-level IFN-γ in TIF showed decreased stem-like properties (Fig. 1J). We also found similar results in tumor tissues from ESCC, CRC and HCC patients, whose low-level border of IFN-γ correlated with highest stem-like properties was 0.072 ng/mL, 0.152 ng/mL and 0.118 ng/mL respectively (Fig. S2B-D). Therefore, we preliminarily surmise that the low-level IFN-γ may serve as a key determinant inducing tumor stemness.

### Low-dose IFN-γ augments the stem-like properties of NSCLC cells

To clarify the role of low-level IFN-γ in inducing tumor stemness, NSCLC cell lines, A549 and H460 cells, were treated with different doses of recombinant human IFN-γ for 1-6 day(s). As expected, the apoptosis rate of the cells progressively increased with an increasing IFN-γ dose and time (Fig. 2A). Strikingly, we observed that the frequency of Annexin V^−^CD133^+^ cells increased over time after treatment with a relatively low dose of IFN-γ (≤0.2ng/mL). In consistent with the optimal IFN-γ concentration correlated with highest stem-like properties in NSCLC TIF (0.110 ng/mL), the optimal concertation of IFN-γ inducing CD133^+^ cells *in vitro* was 0.1 ng/mL. However, the percentage of CD133^+^ cells turned to decrease over time after treated with a relatively high dose of IFN-γ (≥100 ng/mL) (Fig. 2B). Therefore, we speculated that low-dose and long-term IFN-γ stimulation might enhance the stem-like properties of NSCLC cells. To validate this conception, we next sorted the Annexin V^−^ NSCLC cells treated with either low-dose (0.1 ng/mL) or high-dose (100 ng/mL) IFN-γ to examine their sphere-formation ability. Significantly increased sphere number was observed in the low-dose IFN-γ treatment group compared with the control and the high-dose IFN-γ treatment groups (Fig. 2C). Similar results were also found in the expression of CSC markers (Fig. 2D) and CSC signature genes (Fig. 2E). Since high tumorigenicity is a key characteristic of CSCs, we additionally explored the effect of low dose of IFN-γ on tumor growth in NSCLC xenograft models. A549 cells stably expressing luciferase were injected subcutaneously into BABL/c nude mice and then intratumorally treated with PBS or different doses of IFN-γ. Interestingly, we found that both tumor volumes (Fig. 2F, G) and the frequency of CD133^+^ tumor cells in xenografts (Fig. 2H) were dramatically increased following low-dose (0.1 μg/day) but decreased following high-dose (10 μg/day) IFN-γ administration. Taken together, these results suggest that low dose of IFN-γ facilitates the stem-like properties of NSCLC cells both *in vitro* and *in vivo*.

**Figure 2.**
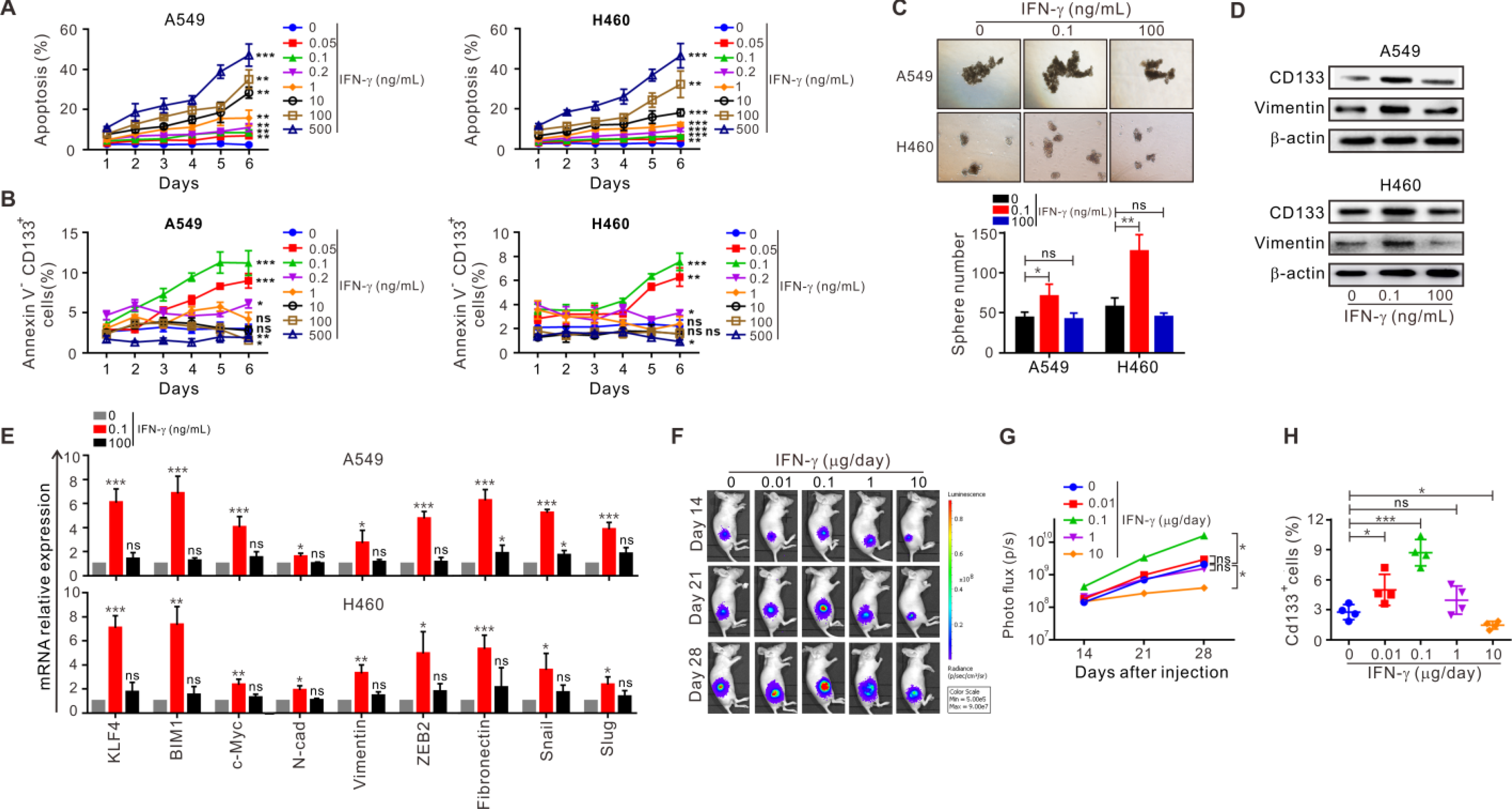
Low-dose IFN-γ augments the stem-like properties of NSCLC cells. A549 and H460 cells were treated with different doses of recombinant human IFN-γ for 1-6 day(s). **(A)** The frequency of apoptotic A549 (left) and H460 (right) cells labeled with Annexin V^+^PI^−^ or Annexin V^+^PI^+^ was examined by flow cytometry. **(B)** The frequency of Annexin V^−^CD133^+^ A549 (left) and H460 (right) cells was examined by flow cytometry. **(C-E)** Annexin V^−^ A549 and H460 cells treated with either low (0.1 ng/mL) or high (100 ng/mL) dose of IFN-γ for 6 days were then sorted using flow cytometry. **(C)** The sphere-formation assay. **(D)** Expression of CD133 and Vimentin in A549 (upper) and H460 (bottom) cells was detected by western blotting. **(E)** The expression of CSC signature genes in A549 (left) and H460 (right) cells was detected by RT-PCR. **(F-H)** A549 cells stably expressing luciferase were injected subcutaneously into nude mice. PBS or different doses of IFN-γ were intratumorally injected from day 14 to 28. **(F, G)** Tumor volumes were measured at day 14, 21 and 28 by *in vivo* imaging system. **(H)** The frequency of ICAM1^+^CD133^+^ tumor cells in mouse-bearing xenografts were detected by flow cytometry. The results are representative of three independent experiments. \**P*<0.05, \*\**P*<0.01, \*\*\**P*<0.001.

### ICAM1 is crucial for the increased stem-like properties of NSCLC cells induced by low-dose IFN-γ

To understand the molecular mechanism underlying low dose of IFN-γ inducing the stem-like properties in NSCLC cells, we searched the IFN-γ signaling downstream genes using the GCBI website (https://www.gcbi.com.cn/) and evaluated the different expression levels of these genes by real-time reverse transcription-PCR (RT-PCR) in A549 and H460 cells treated with or without low dose of IFN-γ. Most of these genes, including IFNGR1 and IFNGR2, were significantly upregulated (Fig. 3A). Among them, ICAM1 showed the most obvious gene upregulation in response to low dose of IFN-γ (Fig. 3A), which was further confirmed by flow cytometry (Fig. 3B). After neutralization with anti-IFN-γ antibody, ICAM1(Fig. S3A) expression was efficiently blocked, as well as IFNGR1 and IFNGR2 (Fig. S3B, C). ICAM1 is a transmembrane molecule that stabilizes cell-cell interactions and enhances transendothelial transmigration(Kanters et al, 2008; Millan et al, 2006). In NSCLC cells, we observed an obvious upregulation of ICAM1 in Calu-3 cells (a highly metastatic NSCLC cell line) and CSCs generated by non-adhesive culture system (Fig. S4), suggesting a potential role of ICAM1 in mediating NSCLC stemness and metastasis. Moreover, IF colocalization analysis demonstrated that ICAM1 was co-expressed with CD133 and Vimentin in A549 cells, which was dramatically increased by low-dose IFN-γ (Fig. 3C). Based on these results, we hypothesized that ICAM1 might be essential for low-dose IFN-γ inducing the stem-like properties in NSCLC cells.

**Figure 3.**
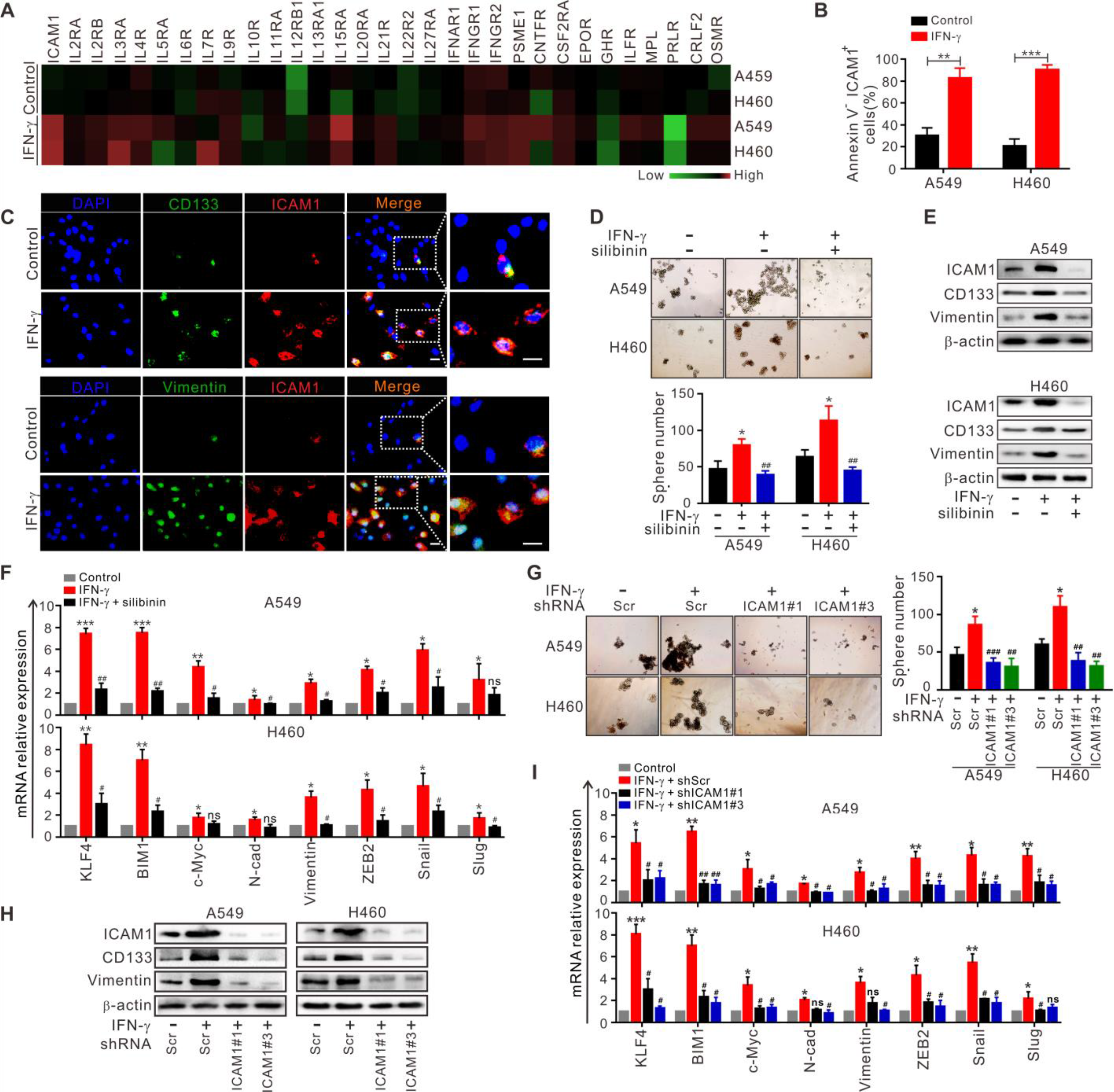
ICAM1 mediates low-dose IFN-γ-induced stem-like properties of NSCLC cells. **(A)** The expression of IFN-γ signaling downstream genes in A549 and H460 cells treated with or without low-dose IFN-γ (0.1 ng/mL) for 6 days was detected by RT-PCR. **(B)** The frequency of Annexin V^−^ICAM1^+^ cells was detected by flow cytometry. **(C)** IF staining for the co-localization of ICAM1 (red) with CD133 (green, upper) or Vimentin (green, bottom) in A549 cells treated with or without low-dose IFN-γ (0.1 ng/mL) for 6 days was performed, and the colocalization is indicated in yellow. DAPI, blue. Scale bars, 20 μm. **(D-F)** A549 and H460 cells were treated with or without silibinin (100 μM) following low-dose IFN-γ (0.1 ng/mL) treatment. Sillibnin and IFN-γ treatment were replicated at day 3 and 5, totally 6 days’ treatment. **(D)** The sphere formation assay in A549 (upper) and H460 (bottom) cells. **(E)** The expression of CD133 and Vimentin in A549 (upper) and H460 (bottom) cells was detected by western blotting. **(F)** The expression of CSC signature genes in A549 (upper) and H460 (bottom) cells was detected by RT-PCR. **(G-I)** A549 and H460 cells were transfected with shScramble, shICAM1#1 or shICAM1#3 following low-dose IFN-γ (0.1 ng/mL) treatment for 6 days. **(G)** The sphere-formation assay. **(H)** The expression of CD133 and Vimentin in A549 (left) and H460 (right) cells was examined by western blotting. **(I)** RT-PCR was used to detect the mRNA expression of CSC signature genes in A549 (upper) and H460 (bottom) cells. Scr, Scramble. The results are representative of three independent experiments.

To test this hypothesis, silibinin, a pharmacological inhibitor of ICAM1, was added to A549 and H460 cells following treatment with low dose of IFN-γ. Interestingly, we found silibinin partially abrogated the sphere-formation ability (Fig. 3D) and the expression of CSC markers (Fig. 3E) and CSC signature genes (Fig. 3F) in NSCLC cells treated with low dose of IFN-γ. Next, we stably knocked down ICAM1 expression in A549 and H460 cells with shICAM1s viruses (Fig. 3H). As expected, ICAM1 knockdown remarkably reduced the stem-like properties induced by low-dose IFN-γ (Fig. 3G-I). Taken together, these results indicate that low dose of IFN-γ drives ICAM1 expression to mediate the stem-like properties of NSCLC cells.

### Activation of the PI3K-Akt-Notch1 pathway by ICAM1 in NSCLC cells is required for low-dose IFN-γ-induced stem-like properties

Next, we explored the signaling pathways involved in the stem-like properties driven by low dose of IFN-γ. We chose several CSC-relevant signaling activation inhibitors, including PI3K, Akt, Notch1, STAT3, p38 and ERK1/2 inhibitors, to pretreat A549 and H460 cells following low-dose IFN-γ treatment. Intriguingly, the RT-PCR results showed that none of the inhibitors had an obvious effect on ICAM1 expression compared with DMSO, but PF04691502, AZD5363 and LY3039478 partially reversed the elevated stem-like properties of A549 and H460 cells (Fig. 4A, B). Similar results were also observed in the flow cytometry and western blot analyses, which demonstrated that PF04691502, AZD5363 and LY3039478 did not affect ICAM1 expression but dramatically blocked the increased expression of CSC markers compared with DMSO in A549 and H460 cells treated with low-dose IFN-γ (Fig. 4C, D). Therefore, we hypothesized that the PI3K-Akt and Notch1 pathways might act as the downstream signaling pathways of ICAM1 to participate in low dose of IFN-γ inducing the stem-like properties in NSCLC cells. The next western blot analysis demonstrated a significantly upregulated activation of Akt and Notch1 after low-dose IFN-γ treatment in contrast to the control, whereas inhibition of ICAM1 by silibinin or ICAM1 shRNAs could efficiently impair their activation (Fig. 4E, F), indicating that ICAM1 activates the PI3K-Akt and Notch1 pathways to mediate the stem-like properties of NSCLC cells treated with low-dose IFN-γ.

**Figure 4.**
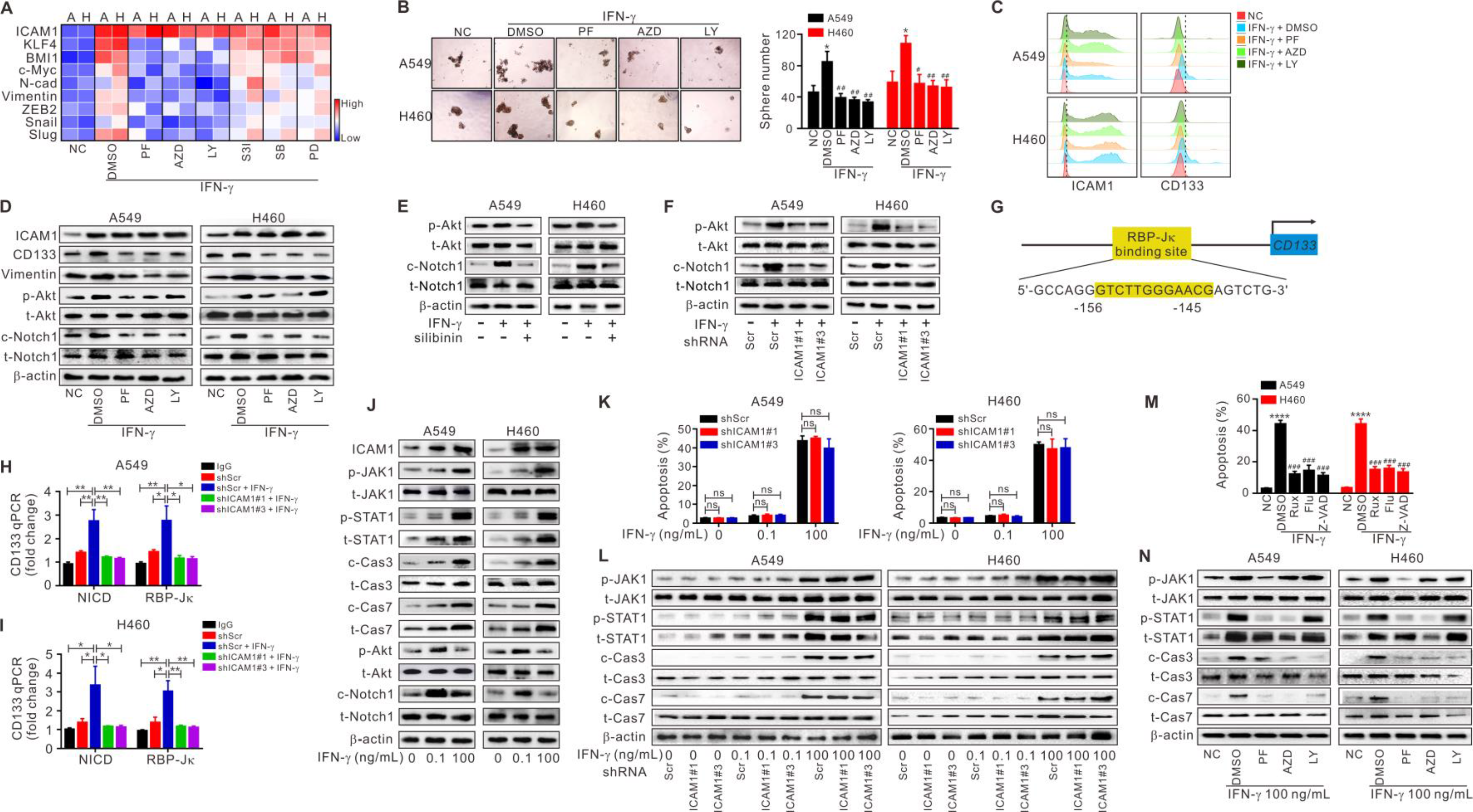
Activation of the ICAM1-PI3K-Akt-Notch1 axis and JAK1-STAT1-caspase pathway are responsible for low-dose-induced stemness and high-dose-mediated apoptosis in IFN-γ treatment system, respectively. **(A-D, M-N)** A549 and H460 cells were pretreated with DMSO, PF04691502 (1 μM), AZD5363 (10 μM), LY3039478 (50 μM), S3I-201 (20 μM), SB203580 (10 μM), PD98059 (20 μM), Ruxolitinib (1μM), Fludarabine (1 μM) or Z-VAD-FMK (40 μM) following low (0.1 ng/mL) or high (100 ng/mL)-dose IFN-γ treatment. Inhibition treatment and IFN-γ treatment were replicated at day 3 and 5, totally 6 days’ treatment. **(A)** The mRNA expression of ICAM1 and CSC signature genes was tested by RT-PCR. **(B)** The sphere-formation assay. **(C)** Flow cytometry was used to test the expression of ICAM1 (left) and CD133 (right). **(D)** Western blot was used to assess the expression of ICAM1, CD133, Vimentin, phospho-Akt, total-Akt, cleaved-Notch1 and total-Notch1. **(E, F)** The expression of phospho-Akt, cleaved-Notch1 and their total proteins was tested by western blotting. **(G)** Analysis of the *CD133* promoter identified an RBP-Jκ-binding site. **(H, I)** ChIP was performed using IgG, anti-NICD or anti-RBP-Jκ antibody, followed by quantitative RT-PCR. **(J)** The expression of ICAM1, phospho-JAK1, phospho-STAT1, cleaved-caspase3, cleaved-caspase7, phospho-Akt, cleaved-Notch1 and their total proteins was tested by western blotting. **(K, L)** Cells were treated with 0, 0.1 and 100 ng/mL IFN-γ for 6 days. **(K, M)** Flow cytometry was used to test apoptosis. **(I, N)** Western blot analysis was performed to examine the expression of phospho-JAK1, phospho-STAT1, cleaved-caspase3, cleaved-caspase7 and their total proteins. A, A549; H, H460; NC, negative control; Scr, Scramble. The results are representative of three independent experiments. In **B** and **M**, \**P*<0.05, \*\*\*\**P*<0.0001 versus negative control group; ^#^*P*<0.05, ^##^*P*<0.01, ^###^*P*<0.001 versus IFN-γ or IFN-γ plus DMSO group. In **H** and **I**, \**P*<0.05; \*\**P*<0.01.

Then, we assessed the activation of the PI3K-Akt and Notch1 pathways in NSCLC cells pretreated with DMSO, PI3K, Akt and Notch1 signaling activation inhibitors following low-dose IFN-γ stimulation. As shown in Fig. 4d, compared with DMSO, PF04691502 and AZD5363 pretreatment reduced the activation of both Akt and Notch1, while LY3039478 pretreatment only attenuated Notch1 activation but did not affect Akt activation. These results suggest that Notch1 is the downstream pathway of the PI3K-Akt pathway in NSCLC cells treated with low-dose IFN-γ.

Upon activation, the Notch1 receptor is cleaved and subsequently releases the Notch1 intracellular domain (N1ICD), which migrates into the nucleus and forms a complex with the nuclear proteins of the RBP-Jκ family. RBP-Jκ acts as a transcriptional activator to activate the expression of target genes after forming a complex with N1ICD(Ohtsuka et al, 1999). We next used the PROMO website to predicate putative RBP-Jκ binding sites in the *CD133* promoter region and identified an RBP-Jκ binding site in the −156 to −145 region (Fig. 4G). Chromatin immunoprecipitation assay (ChIP) assays were performed to validate whether the RBP-Jκ-NICD complex bound to the putative site. As shown in Fig. 4H and I, RBP-Jκ and NICD bound directly to the putative RBP-Jκ-binding site in the *CD133* promoter region in A549 and H460 cells, which showed an increase in shScramble cells treated with low-dose IFN-γ but a decrease in shICAM#1 and shICAM1#3 cells treated with low-dose IFN-γ compared with shScramble cells. These data suggest that Notch1 directly regulates *CD133* transcription in NSCLC cells.

### High-dose IFN-γ contributes to apoptosis through activation of the JAK1-STAT1-caspase signaling pathway in NSCLC cells

Above data revealed that low dose of IFN-γ promoted the stem-like properties of NSCLC cells through activation of the ICAM1-PI3K-Akt-Notch1 axis. However, high dose of IFN-γ was found to induce apoptosis and decrease the stem-like properties of NSCLC cells, as shown in Fig. 2, which prompted our interest in the signaling pathways involved in the high-dose IFN-γ treatment system. The JAK1-STAT1-caspase pathway has been previously described as the key downstream pathway of IFN-γ-mediated apoptosis(Platanias, 2005), and our results indicated that the ICAM1-PI3K-Akt-Notch1 axis was the key signaling cascade in IFN-γ-mediated stem-like properties. Therefore, we next examined the activation of the JAK1-STAT1-caspase pathway and ICAM1-PI3K-Akt-Notch1 axis in NSCLC cells treated with either low-dose or high-dose IFN-γ, respectively. The results showed that cells treated with high-dose IFN-γ exhibited much higher activation of JAK1, STAT1, caspase3 and caspase7 than those treated with low-dose IFN-γ (Fig. 4J). Surprisingly, we further observed a significant decrease in the activation of Akt and Notch1 in cells treated with high-dose IFN-γ compared to those treated with low-dose of IFN-γ although both high-dose and low-dose IFN-γ were able to induce high levels of ICAM1 expression (Fig. 4J). Thus, high-dose IFN-γ promoted apoptosis of NSCLC cells through the JAK1-STAT1-caspase pathway, whereas high-dose IFN-γ-induced ICAM1 was insufficient to endow the stem-like properties of NSCLC cells through the PI3K-Akt-Notch1 signaling pathway.

Next, we wondered whether ICAM1 played a role in high-dose IFN-γ-mediated apoptosis. After treatment with different doses of IFN-γ, we found that ICAM1 knockdown had no influence on apoptosis in A549 and H460 cells (Fig. 4K), while activation of JAK1, STAT1, caspase3 and caspase7 was increased with the increase of IFN-γ dose (Fig. 4L). Moreover, inhibition of JAK1 (Ruxolitinib), STAT1 (Fludarabine) and caspase (Z-VAD-FMK) significantly attenuated high-dose IFN-γ-mediated apoptosis (Fig. 4M). Western blot analysis showed that phospho-JAK1 expression was only inhibited by Ruxolitinib and phosoho-STAT1 expression was inhibited by both Ruxolitinib and Fludarabine, while caspase 3 and 7 were inhibited by Ruxolitinib, Fludarabine and Z-VAD-FMK (Fig. 4N). These data indicate that high-dose IFN-γ promotes apoptosis of NSCLC cells by activating the JAK1-STAT1-caspase signaling pathway in an ICAM1-independent manner.

### ICAM1 knockdown reverses tumor growth induced by low-dose IFN-γ in xenograft model of NSCLC

To identify whether ICAM1 played a role in low-dose IFN-γ-induced tumor growth, A549 cells stably expressing luciferase were injected subcutaneously into BALB/c nude mice. The mice were treated intratumorally with PBS or low-dose IFN-γ, silibinin was orally administered. We found that the enhanced tumor growth induced by low-dose IFN-γ was efficiently blocked by silibinin (Fig. S5A, B). Silibinin also dramatically reduced the high frequency of ICAM1^+^CD133^+^ tumor cells in xenografts treated with low-dose IFN-γ (Fig. S5C).

In Fig. 3F, G, we had confirmed that low dose of IFN-γ promoted NSCLC growth via ICAM1 in BABL/c nude mice. Next, NOD-SCID mice were used as another model to evaluate the protumor effect of low-dose IFN-γ and the role of ICAM1 during this process. Luciferase-shScramble-infected, luciferase-shICAM1#1-infected and luciferase-shICAM1#3-infected cell lines were stably established and injected subcutaneously into NOD-SCID mice. The mice in each group were further intratumorally treated with PBS or low-dose IFN-γ. We found tumor growth of shICAM1#1 cell-derived and shICAM1#3 cell-derived xenografts was significantly reduced compared with shScramble cell-derived ones (Fig. 5A, B). Moreover, a remarkable increase of tumor growth was detected in shcaramble cell-derived xenografts after treatment with low-dose IFN-γ, while shICAM1#1 and shICAM1#3 cell-derived xenografts did not exhibit obvious differences after low-dose IFN-γ treatment (Fig. 5A, B). Besides, low-dose IFN-γ treatment greatly enhanced the frequency of ICAM1^+^CD133^+^ tumor cells in shScramble cell-derived xenografts but had no significant effect on those in shICAM1#1 and shICAM1#3 cell-derived xenografts (Fig. 5C). Similar results were obtained for the expression of ICAM1, CD133 and Vimentin by IHC staining (Fig. 5D-G). Taken together, these data indicate that ICAM1 is essential for low-dose IFN-γ-mediated tumor growth in immunodeficient mice.

**Figure 5.**
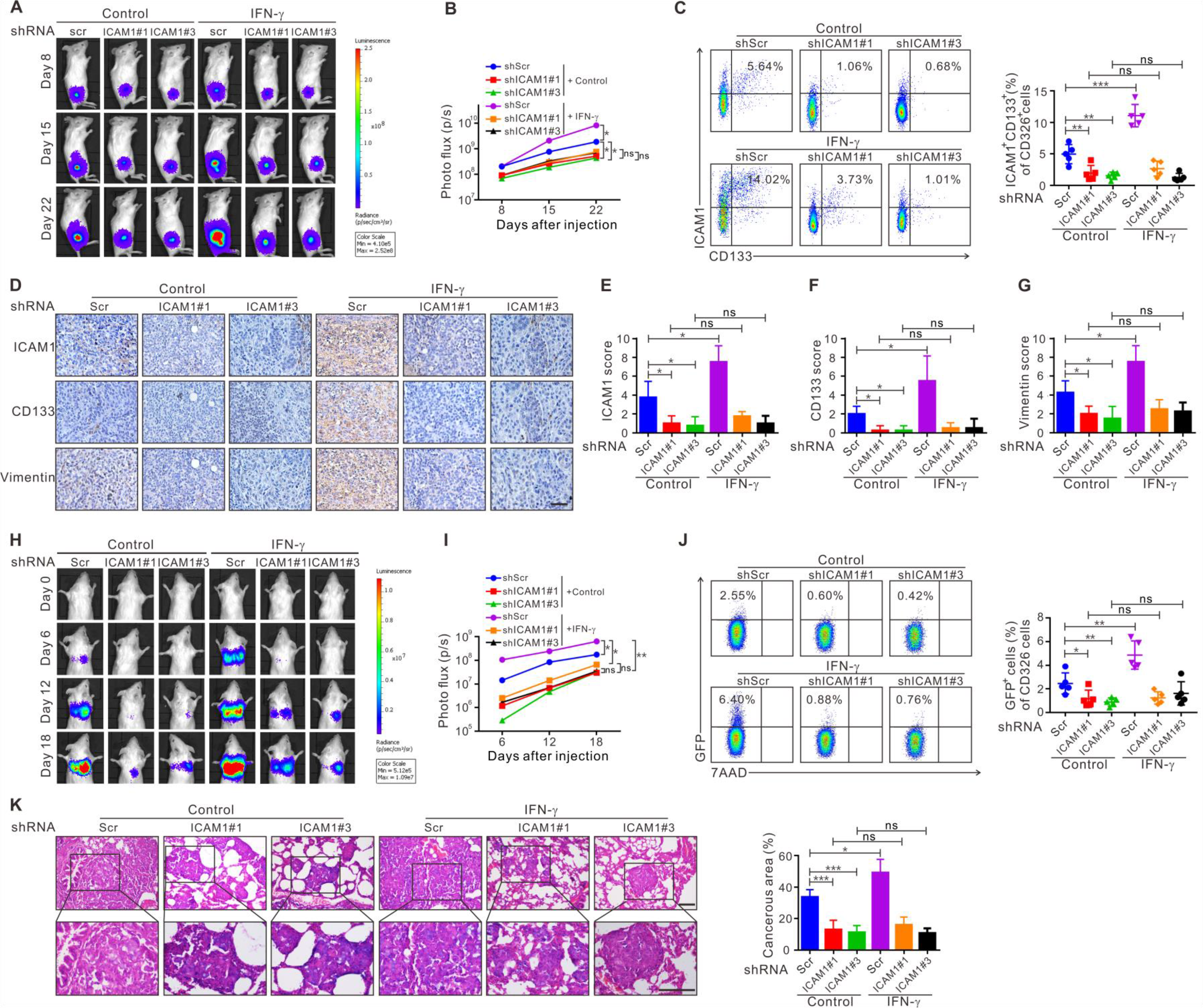
Low-dose IFN-γ promotes tumor growth and lung metastasis of NSCLC in an ICAM1-dependent manner. **(A, B)** Tumor growth was measured at day 14, 21 and 28 after cell injection using in vivo imaging system. **(C)** The frequency of ICAM1^+^CD133^+^ tumor cells of CD326^+^ cells in mice-bearing xenografts were detected using flow cytometry. **(D)** Consecutive sections of mice-bearing xenografts were used to analyze the co-expression levels of ICAM1, CD133 and Vimentin by IHC staining. Scale bars, 50μm. **(E-G**) Scores of ICAM1, CD133 and Vimentin in mice-bearing xenografts are shown as a statistical graph. **(H, I)** Metastasis was evaluated using an *in vivo* imaging system at day 0, 6, 12 and 18 after injection. The lungs of the mice in each group were harvested at day 22 to analyze the frequency of GFP^+^ NSCLC cells of CD326^+^ cells by flow cytometry. **(J)** and the metastatic nodules in the lungs by hematoxylin-eosin staining. **(K)**. Scale bars, 100 μm. Scr, Scramble. The results are representative of three independent experiments. \**P*<0.05, \*\**P*<0.01, \*\*\**P*<0.001.

### Low-dose IFN-γ facilitates NSCLC cell metastatic growth in murine lungs via ICAM1

Apart from the high sphere-formation ability and high tumorigenicity, CSCs can also form a hierarchy of stem-like and differentiated tumor cells to initiate metastatic growth(Plaks et al, 2015). Therefore, we investigated the effect of low-dose IFN-γ on the metastatic growth of NSCLC cells and the role of ICAM1 in this process. Before intravenous injection into the lateral tail vein of NOD-SCID mice, luciferase-shScramble-infected, luciferase-shICAM1#1-infected and luciferase-shICAM1#3-infected cells were pretreated with PBS or low-dose IFN-γ for 6 days *in vitro*. Metastasis was evaluated by luminescence imaging at day 0, 6, 12 and 18 after injection. We observed that luciferase-shScramble-infected cells pretreated with low-dose IFN-γ generated larger lung metastatic nodules than luciferase-shScramble-infected cells pretreated with PBS, whereas luciferase-shICAM1#1- and shICAM1#3-infected cells pretreated with low-dose IFN-γ did not show significant difference compared with those pretreated with PBS (Fig. 5H, I). In the PBS-pretreated groups, mice injected with luciferase-shICAM1#1- and shICAM1#3-infected cells had smaller lung metastatic nodules than those injected with luciferase-shScramble-infected cells (Fig. 5H, I). Consistently, similar results were also obtained in the frequency of green fluorescent protein (GFP)^+^ NSCLC cells (Fig. 5J) and the percentage of metastatic cancerous areas in the lungs of mice (Fig. 5K). Together, these results suggest that low-dose IFN-γ greatly enhances lung metastasis of NSCLC cell *in vivo*, which is mediated by ICAM1.

### The expression level of ICAM1 is significantly upregulated by CD133^+^ tumor cells and positively correlated with a poor prognosis in NSCLC patients

Subsequently, we detected the expression level of ICAM1 in clinical samples. By flow cytometry analysis, we observed higher ICAM1 expression by tumor cells from tumor tissues than epithelial cells from adjacent normal tissues in patients with NSCLC (Fig. 6A). Moreover, a dramatically elevated frequency of ICAM1^+^ cells in CD133^+^ tumor cells was observed compared with CD133^−^ tumor cells (Fig. 6B). Data from the TCGA database revealed a close association of ICAM1 with CSC related genes at the mRNA level in 1016 NSCLC tissues (Fig. S6A-G). Consistently, the IHC staining results demonstrated that patients with high-level ICAM1 expression had higher CD133 (Fig. 6C) and Vimentin (Fig. 6D) expression than those with low-level ICAM1 expression. Strongly positive correlation of ICAM1 with CD133 (Fig. 6E) and Vimentin (Fig. 6F) was also observed in these tissues. Moreover, co-expression of ICAM1 with CD133 and Vimentin was found in NSCLC tissues, and the co-expression levels in chemoresistant patients were significantly higher than those in chemosensitive patients (Fig. S6H). These data suggest that ICAM1 may be a malignant signature of NSCLC cells.

**Figure 6.**
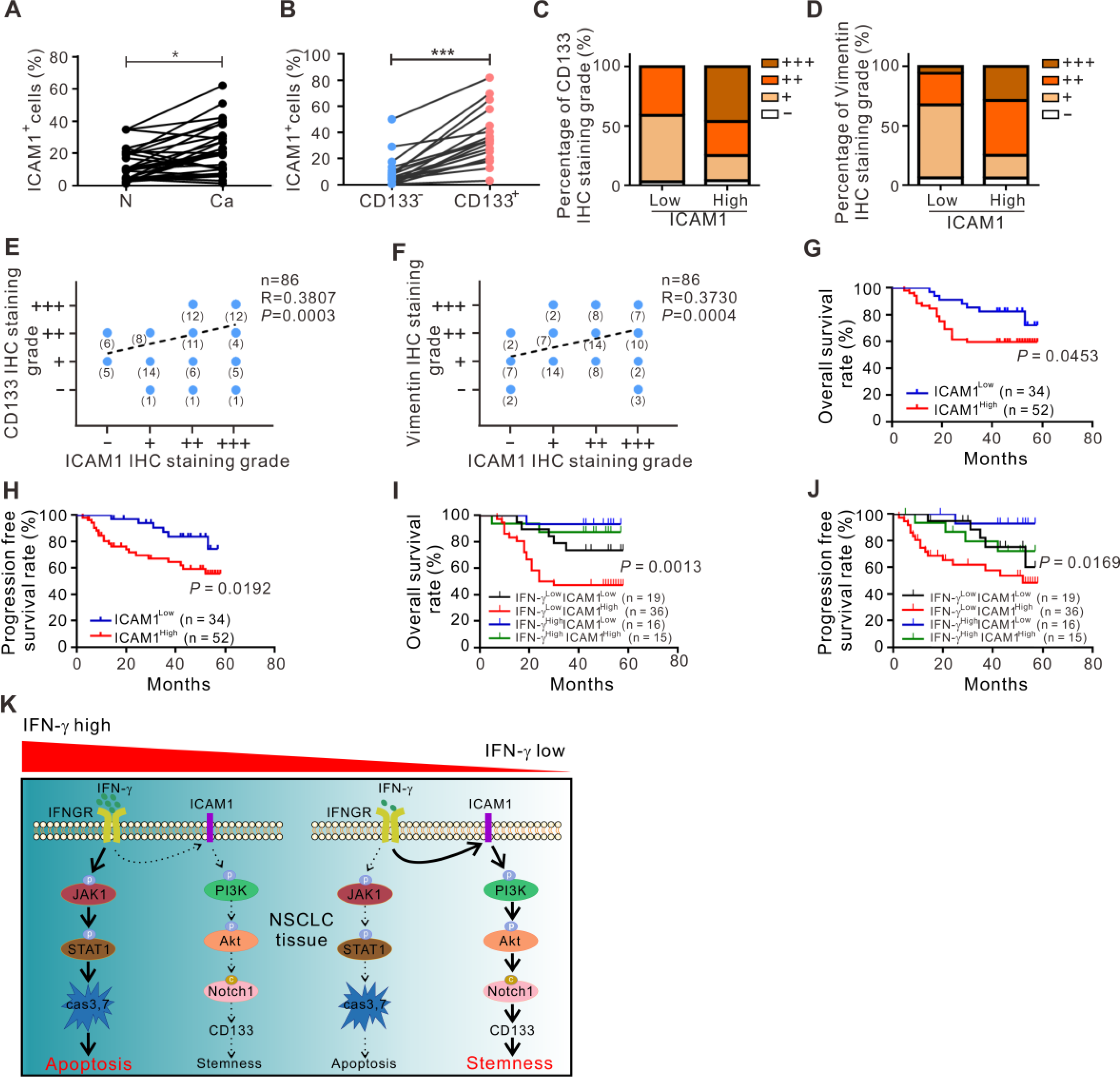
ICAM1 is significantly upregulated by CD133^+^ tumor cells and strongly correlated with a poor prognosis in NSCLC patients. **(A)** The frequency of ICAM1^+^ cells in CD326^+^ cells in tumor tissues and adjacent normal tissues were analyzed by flow cytometry (n=35). **(B)** The frequency of ICAM1^+^ cells in CD326^+^CD133^−^ and CD326^+^CD133^+^ cells were analyzed by flow cytometry in NSCLC tissues (n=35). **(C, D)** The actual distribution of IHC results between ICAM1 and CD133 or Vimentin expression in 86 NSCLC tissues. **(E, F)** Spearman’s correlation analysis of the correlation between ICAM1 and CD133 or Vimentin in 86 NSCLC specimens was performed according to the IHC staining results. **(G-J)** Kaplan-Meier analysis of overall survival and progression-free survival in the 86 NSCLC patients according to ICAM1 or both IFN-γ and ICAM1 expression assessed by IHC staining. **(K)** Schematic diagram. p, phosphorylation; c, cleaved; cas, caspase. Log-rank tests were used in **G-J**. N, normal tissue; Ca, cancer tissue. The results are representative of three independent experiments. \**P*<0.05, \*\*\**P*<0.0001.

Based on the IHC staining of ICAM1, a significant correlation was found between ICAM1 expression and TNM stage, recurrence, bone metastasis, epidermal growth factor receptor (EGFR) mutation and chemoresistance (Table 2). Patients with high-level ICAM1 had a worse OS (Fig. 6G) and PFS (Fig. 6H). Data from TCGA also showed that high-level ICAM1 was strongly associated with a poor PFS in NSCLC patients (Fig. S5I). These data suggest that ICAM1 is strongly associated with prognosis in NSCLC patients. In addition, we further stratified the 86 NSCLC tissues into four groups according to the expression levels of IFN-γ and ICAM1 analyzed by IHC staining: IFN-γ^low^ICAM1^low^, IFN-γ^low^ICAM1^high^, IFN-γ^high^ICAM1^low^ and IFN-γ^high^ICAM1^high^. Surprisingly, we found that patients with IFN-γ^high^ICAM1^high^ expression showed a better OS (Fig. 6I) and PFS (Fig. 6J) than those with IFN-γ^low^ICAM1^high^ expression, which supported the view that ICAM1 might mediate tumor progression or even stemness only in an IFN-γ-low TME rather than in an IFN-γ-high TME.

**Table 2.** Association of ICAM1 expression with clinicopathological features in patients with NSCLC.

| Characteristic | Total | ICAM1 expression (n=86) | | $\chi^2$ | P-value |
| --- | --- | --- | --- | --- | --- |
|  |  | Low<br>n=34(40%) | High<br>n=52(60%) |  |  |
| Age, years |  |  |  |  |  |
| $\geq 55$ | 61 | 23 | 38 | 0.294 | 0.5877 |
| <55 | 25 | 11 | 14 |  |  |
| Gender |  |  |  |  |  |
| Male | 60 | 27 | 33 | 2.48 | 0.1153 |
| Female | 26 | 7 | 19 |  |  |
| Smoking History |  |  |  |  |  |
| Yes | 45 | 17 | 28 | 0.1219 | 0.7270 |
| No | 41 | 17 | 24 |  |  |
| Differentiation |  |  |  |  |  |
| Well/Moderate | 66 | 27 | 39 | 0.2242 | 0.6359 |
| Poor | 20 | 7 | 13 |  |  |
| Histology |  |  |  |  |  |
| Adenocarcinoma | 47 | 13 | 34 | 3.423 | **0.0134\*** |
| Squamous cell carcinoma | 39 | 21 | 18 |  |  |
| TNM stage |  |  |  |  |  |
| I-II | 63 | 29 | 34 | 4.159 | **0.0414\*** |
| III-IV | 23 | 5 | 18 |  |  |
| Recurrence |  |  |  |  |  |
| Yes | 14 | 2 | 12 | 4.46 | **0.0347\*** |
| No | 72 | 32 | 40 |  |  |
| Lymph node metastasis |  |  |  |  |  |
| Yes | 29 | 12 | 17 | 0.06227 | 0.8029 |
| No | 57 | 22 | 35 |  |  |
| Bone metastasis |  |  |  |  |  |
| Yes | 13 | 1 | 12 | 6.496 | **0.0108\*** |
| No | 73 | 33 | 40 |  |  |
| Brain metastasis |  |  |  |  |  |
| Yes | 7 | 3 | 4 | 0.05714 | 0.8111 |
| No | 81 | 31 | 50 |  |  |
| EGFR mutation |  |  |  |  |  |
| Yes | 34 | 9 | 25 | 4.015 | **0.0451\*** |
| No | 52 | 25 | 27 |  |  |
| Chemoresistance |  |  |  |  |  |
| Yes | 20 | 3 | 17 | 6.562 | **0.0104\*** |
| No | 66 | 31 | 35 |  |  |

EGFR, epidermal growth factor receptor.
ICAM1, intercellular cell adhesion molecule-1.
NSCLC, non-small cell lung cancer.
\* $P < 0.05$ .

## Discussion

Emerging evidences has reported that IFN-γ may involve in tumor progression, but the exact mechanism was not fully understood. In this study, we firstly identified the dose-dependent effect of IFN-γ on tumor stemness in NSCLC. Low-dose IFN-γ induced tumor stemness via the ICAM1-PI3K-Akt-Notch1 axis, whereas high-dose IFN-γ mainly mediated cell apoptosis through the JAK1-STAT1-caspase pathway (Fig. 6K).

IFN-γ has a controversial role in regulating antitumor immunity. As there are few immunosuppressive factors in the TME in the early stage of solid tumor growth, effector T and NK cells can efficiently recognize tumor cells and subsequently produce high levels of IFN-γ, resulting in the apoptosis of tumor cells. However, with tumor progression, T and NK cells become dysfunctional in the late stage(Wherry & Kurachi, 2015), eventually generating an IFN-γ-low TME. In addition, there is a reduction of T and NK cell infiltration in the center of cancer nest as well as those “cold tumors”, which may also form an IFN-γ-low niche, lacking effective immune infiltration. Low-level IFN-γ is not sufficient to induce the apoptosis of a large number of tumor cells, which in turn accelerates tumor development through various mechanisms, such as the mediation of immunosuppression (Gato-Canas et al, 2017; Katz et al, 2008; Spranger et al, 2013), maintenance of CSCs dormancy(Liu et al, 2017) and induction of metastatic CSC generation(Chen et al, 2011). In this study, we clearly demonstrated that low-dose IFN-γ endowed stem-like properties of tumor cells through the ICAM1-PI3K-Akt-Notch1 axis, while high-dose IFN-γ mainly exerted antitumor effect via the JAK1-STAT1-caspase pathway in NSCLC. According to these findings, we infer that the regulation of low-level IFN-γ on tumor stemness *in vivo* may be a spatial and temporal effect, which only occur in the time of in late-stage or in the sites lacking effective immune infiltration. Consistently, the function of low-level IFN-γ in maintaining or promoting tumor stemness has also been reported in certain tumor models(Chen et al, 2011; Liu et al, 2017). However, since it is currently very difficult to precisely control the level of IFN-γ in immunocompetent mice due to unknown IFN-γ production in many cells and tissues, we only verified our results *in vivo* by using immunodeficient mouse models in this study. Moreover, IFN-γ induces not only apoptosis or the stemness of tumor cells but also strongly upregulates PD-L1 expression on tumor cells, activates immunosuppressive myeloid-derived suppressor cells by inducing iNOS. All these points will be taken into account in *in vivo* experiments in our further study.

Clinically, the adoptive transfer of receptor-engineered T cells and immune checkpoints blockade are still limited in treating some solid tumors(Ping et al, 2018; Zou, 2018). It was reported that low-dose IFN-γ generated at the tumor site increases the risk of tumor metastasis during immunotherapy(Gong et al, 2008; Kelly et al, 1991). A recent study has reported that tumor stemness might cause PD-1/PD-L1 blockade resistance(Zou et al, 2016). In our view, the low-level IFN-γ generated by dysfunctional T or NK cells in the TME may lead to treatment failure of immunotherapy by inducing tumor stemness. The dose of IFN-γ delivered to the tumor site should be high when using IFN-γ-based cancer immunotherapy. Accordingly, for adoptive T-cell therapy, a key issue is to ensure the effector function of T cells after reaching the tumor site, particularly when treating advanced solid tumors.

In this study, we found that ICAM1 merely mediated tumor stemness in the IFN-γ-low rather than the IFN-γ-high TME, and better survival was also observed in IFN-γ^high^ICAM1^high^ patients that that in IFN-γ^low^ICAM1^high^ patients. It may be necessary to evaluate the levels of IFN-γ in the TME when using ICAM1-targeting therapy, as patients with low-IFN-γ levels may be more sensitive. However, ICAM1 was expressed on not only tumor cells but also a variety of stroma cells, such as endothelial cells. It cannot be directly used as a specific target. ICAM1 targeting therapy may be applied by using nanoparticles encapsulating ICAM1 small molecular inhibitor, such as silibinin, which can specifically reach tumor sites.

In summary, low-dose IFN-γ preferably induced tumor stemness via the ICAM1-PI3K-Akt-Notch1 axis, whereas high-dose IFN-γ mainly mediated cell apoptosis through the JAK1-STAT1-caspase pathway in NSCLC. Inhibition of ICAM1 efficiently blocked the stem-like properties of tumor cells induced by low-dose IFN-γ *in vitro* and *in vivo*. Our findings firstly revealed the dose-dependent effect of IFN-γ in inducing tumor stemness and clearly elucidated the distinct molecular mechanisms activated by IFN-γ in a dose-dependent manner, providing insights into cancer progression and treatment, particularly patients with low-level IFN-γ expression in the TME.

## Materials and Methods

### Study approval

All patients provided written informed consent according to the consent form of the Ethics Committee of the first affiliated hospital of Zhengzhou University (Research-2015-LW-108). All animal procedures were carried out according to the Guide for the Care and Use of Laboratory Animals and were approved by the Institutional Animal Care and Use committee of the First Affiliated Hospital of Zhengzhou University (Approval No.00015613).

### Patients and tumor samples

For IHC and IF staining, a total of 86 NSCLC tissues were collected from the First Affiliated Hospital of Zhengzhou University from 2012 to 2014. All patients received four cycles of cisplatin-based chemotherapy after surgery without receiving any therapeutic intervention before surgery. For flow cytometry, 35 pairs of NSCLC tissues and their adjacent normal tissues were freshly obtained from the First Affiliated Hospital of Zhengzhou University from January to May 2017. For isolation of interstitial fluid, 9 fresh NSCLC tissues, 9 fresh ESCC tissues, 11 fresh CRC tissues and 13 fresh HCC tissues were collected from the First Affiliated Hospital of Zhengzhou University. The methods for sample collections in the study were approved by the Ethics Committee of the First Affiliated Hospital of Zhengzhou University and confirmed by pathological analysis. The clinicopathologic characteristics, including gender, age, stage, smoking history, differentiation, histological grade, TNM stage, recurrence, lymph node metastasis, bone metastasis, brain metastasis, EGFR mutation and chemoresistance were evaluated.

### NSCLC cell lines and cell culture

The NSCLC cell lines A549, H460 and Calu-3 were purchased from the Chinese Academy of Sciences Cell Repertoire in Shanghai China and maintained in RPMI-1640 (HyClone, Logan, UT, USA) supplemented with 10% fetal bovine serum (HyClone, Logan, UT, USA), 100 units/mL of penicillin and 100 μg/mL of streptomycin at 37°C and 5% CO_2_ in a humidified incubator. The methods of low or high-dose IFN-γ treatment for 6 days in A549 and H460 cells were as follows: recombinant human IFN-γ (Peprotech, Rocky Hill, NJ, USA) were added and cultured for 2 days, and re-stimulated at day 3 and 5, totally 6 days’ treatment. Inhibition and blockade treatment were as follows: anti-IFN-γ antibody (R&D system lnc., Minneapolis, MN, USA), inhibitors, including silibinin (Sigma, Santa Clara, CA, USA), PF04691502, AZD5363 and LY3039478 (Selleck Chemicals, Houston, TX, USA) were pretreatment for 1h following IFN-γ treatment for 2 days, which were replicated at day 3 and 5, totally 6 days’ treatment.

### Isolation of interstitial fluid from fresh tumor tissues

Fresh tissues were placed on triple-layered 10-μm nylon mesh and spun at <50g for 5min to remove surface liquid. Next, samples were centrifuged at 400g for 10 minutes to isolate the TIF(Eil et al, 2016).

### Transfected cell lines

A549 cells were stably transfected with luciferase expression vector. Stable knockdown of ICAM1 in A549, H460 and A549 stably expressing luciferase cells by short hairpin RNA was achieved using the pGV248-hU6-MCS-Ubiquitin-EGFP-IRES-puromycin vector plasmid purchased from Gene Pharma (Gene Pharma, Shanghai, China). For viral transductions, the lentiviruses were harvested in 293T cells. Then, A549, H460 and A549 stably expressing luciferase cells were infected with shScramble and shICAM1 lentiviruses. After 72 h, the transfected cells were sorted by flow cytometry (Beckman MoFlo XDP, Brea, CA, USA) according to the expression of GFP. The expression of ICAM1 was validated by RT-PCR and western blot analysis. All inserted sequences were verified by DNA sequencing.

### Sphere-formation assay

A549 and H460 cells treated with low or high-dose IFN-γ were sorted. A549 and H460 cells pretreated with DMSO, silibinin (100 μM), PF04691502, AZD5363 and LY3039478 or knocked down ICAM1 with shScramble and shICAM1 RNAs following treatment with low-dose IFN-γ were sorted. A total of 3×10^3^ A549 or H460 cells per well were seeded in ultra-low attachment 24-well plates in DMEM/F12 medium (Sigma) containing B27 supplement (Gibco, Invitrogen Carlsbad, CA, USA), EGF (20 ng/mL; Peprotech) and bFGF (20 ng/mL; Peprotech). After 7 days, spheres with a diameter > 75 μm were counted.

### Animal xenograft models

To explore the effect of low-dose IFN-γ on tumor growth in xenograft models, 20 6-week-old female BALB/c nude mice (Vital River Laboratory Animal Technology Co. Ltd, Beijing, China) were randomly divided into five groups (n=5 each) and hypodermically injected with A549 cells stably expressing luciferase (1 × 10^7^ cells in 100 μL PBS). After two weeks, 100 to 200 mm^3^ tumors were apparent on all mice. Each group was intratumorally treated with PBS or different doses of IFN-γ (0.01, 0.1, 1, 10 μg/day, once per two days) from day 14 to 28. Tumor growth was monitored once per week using an *in vivo* imaging system (PerkinElmer, IVISLumina Series III, USA). Xenografts were harvested at day 28 to analyze the frequency of CD133^+^ tumor cells.

To evaluate the protumor effect of low-dose IFN-γ in NOD-SCID mice and the role of ICAM1 during this process, luciferase-shScramble-infected, luciferase-shICAM1#1-infected and luciferase-shICAM1#3-infected cells were injected subcutaneously into 24 5-week-old female NOD-SCID mice (three groups, n=10 each). After 8 days, the tumors volumes reached 100 to 200 mm^3^ in all mice. At day 8, each group was randomly dived into two groups (a total of six groups, n=5 each). The mice were further intratumorally treated with PBS or low-dose IFN-γ (0.1 μg/day, once per two days) from day 8 to day 22. Tumor growth was measured at day 8, 15 and 22 using an *in vivo* imaging system (PerkinElmer, IVISLumina Series III, USA). All the xenografts were harvested at day 22. Flow cytometry analysis was performed to detect the frequency of ICAM1^+^CD133^+^ tumor cells. IHC staining was performed to examine the expression of ICAM1, CD133 and Vimentin.

In addition, to investigate whether silibinin could reverse the low-dose IFN-γ induced tumor growth *in vivo*, 16 5-week-old female BALB/c nude mice (Vital River Laboratory Animal Technology Co. Ltd, Beijing, China) were further randomly divided into four groups (n=5 each). All the groups were hypodermically injected with A549 cells stably expressing luciferase (1 × 10^7^ cells in 100 μL PBS). Two weeks later, the tumors in each group reached 100 to 200 mm^3^. PBS or low-dose IFN-γ (0.1 μg/day, once per two days) were intratumorally injected from day 14 to 28 concomitantly with oral administration of silibinin (in drinking water, 5 mg/mL, 3-4 mL/mouse/day). Tumor growth was evaluated at day 14, 21 and 28 using an *in vivo* imaging system (PerkinElmer, IVISLumina Series III, USA). Xenografts were harvested to analyze the frequency of ICAM1^+^CD133^+^ tumor cells by flow cytometry. Animal studies were blinded during data measurement.

### In vivo metastatic growth model

Twenty-four 5-week-old female NOD-SCID mice were randomly divided into six groups (n=5 each). Luciferase-shScramble-infected, luciferase-shICAM1#1-infected and luciferase-shICAM1#3-infected cells were pretreated with PBS or low-dose IFN-γ (0.1 ng/mL) *in vitro* for 6 days and then intravenously injected into the lateral tail vein of mice (5 × 10^6^ cells in 100 μL PBS). Metastasis was evaluated using an *in vivo* imaging system at day 0, 6, 12 and 18 after injection. The lungs of the mice in each group were harvested to analyze the frequency of GFP^+^ NSCLC cells by flow cytometry and the metastatic nodules by hematoxylin-eosin staining.

### RNA extraction and quantitative RT-PCR

A549 and H460 cells treated with low or high-dose IFN-γ were sorted. A549 and H460 cells pretreated with DMSO, silibinin, PF04691502, AZD5363 and LY3039478 or with ICAM1 knockdown by shScramble and shICAM1 RNAs following low-dose IFN-γ treatment were sorted. A549 and H460 cells pretreated with DMSO, Ruxolitinib, Fludarabine and Z-VAD-FMK following high-dose IFN-γ treatment were sorted. Total RNA was extracted from NSCLC cells with TRIzol reagent (Invitrogen Corporation, Carlsbad, CA, USA) according to the manufacturer’s instructions. The purity and concentration of the RNA were examined using NanoDrop 2000 (Thermo Fisher Scientific, Waltham, USA). First-strand cDNA was synthesized from 1 μg of total RNA using the PrimeScript™ RT reagent Kit with gDNA Eraser (TaKaRa, Dalian, China). SYBR Premix Ex Taq II (TaKaRa) was used to perform quantitative RT-PCR in Agilent Mx3005P. Each experiment was performed in triplicate. Glyceraldehyde 3-phosphate dehydrogenase (GAPDH) was used as endogenous control for normalization.

### Flow cytometry

Intracellular staining was performed as previously described(Li et al, 2016). Tumor tissues were minced, and the tissue homogenate was then filtered through a 70-μm filter to isolate single cells. A549 and H460 cells pretreated with or without anti-IFN-γ antibody following low-dose IFN-γ stimulation were stained with anti-human CD3, CD4, CD8, CD56, CD326, CD133, ICAM1 or 7-AAD antibodies (BioLegend, San Diego, CA, USA). CD326 was used to identify tumor cells in tumor tissues and epithelial cells in adjacent normal tissues. Additionally, A549 and H460 cells treated with or without IFN-γ were stained with anti-Annexin V, anti-human CD133 and anti-ICAM1 antibodies (BioLegend). A549 and H460 cells pretreated with DMSO, Ruxolitinib, Fludarabine and Z-VAD-FMK following high-dose IFN-γ treatment were stained with anti-Annexin V antibody. Propidium iodide solution was added to examine cell apoptosis. Cells were analyzed by flow cytometry (BD FACSCantoII, USA).

### IHC and IF staining

The protocols used for IHC and IF have been described elsewhere(Qiao et al, 2017). Paraffin-embedded NSCLC tissues or A549 cells treated with or without IFN-γ were examined for the expression of IFN-γ, ICAM1, CD133 or Vimentin. For IHC staining, the following primary antibodies were used: rabbit anti-human IFN-γ (Proteintech Group, Chicago, Illinois, USA, 1:100), rabbit anti-human ICAM1 (Proteintech Group, 1:400), rabbit anti-human CD133 (Thermo Fisher Scientific) or rabbit anti-human Vimentin (Proteintech Group, 1:800). The intensity of the staining was analyzed using the following criteria: 0, negative; 1, low; 2, medium; and 3, high. The extent of staining was scored as 0, 0% stained; 1, 1-25% stained; 2, 26-50% stained; and 3, 51-100% stained. Five random fields were analyzed using a light microscope (Olympus, Tokyo, Japan). The final scores were evaluated by multiplying the scores of the intensity by those of the extent. The samples were divided into four staining grades: 0, −; 1-3, +; 4-6, ++; and 7-9, +++. The following criteria were used to quantify the expression levels of IFN-γ and ICAM1 in NSCLC tissues: low expression, “− or +”; high expression, “++ or +++”. For IF staining, the following primary antibodies were used: mouse anti-human IFN-γ (R&D system, 1:1000), mouse anti-human ICAM1 (Abcam, Cambridge, MA, USA, 1:100), rabbit anti-human CD133 (Thermo Fisher Scientific, 1:1000) or rabbit anti-human Vimentin (Proteintech Group, 1:500). FITC-conjugated anti-rabbit IgG (BioLegend, 1:500) and Cy3-conjugated anti-mouse IgG (BioLegend, 1:500) were used as secondary antibodies. The nuclei were stained with 40,6-diamidino-2-phenylindole (DAPI, Roche, USA, 1:1000). Images were analyzed using a fluorescence microscope (Olympus, IX71, Japan). The IF staining results were quantified using NIH ImageJ software.

### Western blotting

The protocol used for western blot has been described elsewhere(Qiao et al, 2017). A549 and H460 cells treated with low-dose or high-dose IFN-γ were sorted. A549 and H460 cells pretreated with DMSO, silibinin, PF04691502, AZD5363 and LY3039478 or shScramble and shICAM1 RNAs following low-dose IFN-γ treatment were sorted. A549 and H460 cells pretreated with DMSO, Ruxolitinib, Fludarabine and Z-VAD-FMK following high-dose IFN-γ treatment were sorted. The primary antibodies used were as follows: anti-ICAM1 (Abcam, 1:100), anti-CD133 (Proteintech Group, 1:1000), anti-Vimentin (Proteintech Group, 1:500), anti-Akt (Cell Signaling Technology, Boston, MA, USA, 1:1000), anti-phospho-Akt (Cell Signaling Technology, 1:1000), anti-Notch1 (Cell Signaling Technology, 1:1000), anti-cleaved-Notch1 (Cell Signaling Technology, 1:1000), anti-JAK1 (Cell Signaling Technology, 1:1000), anti-phospho-JAK1 (Cell Signaling Technology, 1:1000), anti-STAT1 (Cell Signaling Technology, 1:1000), anti-phospho-STAT1 (Cell Signaling Technology, 1:1000), anti-caspase3 (Cell Signaling Technology, 1:1000), anti-cleaved-caspase3 (Cell Signaling Technology, 1:1000), anti-caspase7 (Cell Signaling Technology, 1:1000), anti-cleaved-caspase7 (Cell Signaling Technology, 1:1000) and anti-β-actin (Cell Signaling Technology, 1:4000).

### ChIP assay

The ChIP assay was performed according to the manufacturer’s instructions (Cell Signaling Technology). Anti-N1ICD (Cell Signaling Technology 1:200) or anti-RBP-Jκ (Cell Signaling Technology, 1:50) antibody was used to immunoprecipitate the chromatin in A549 and H460 cells transfected with shScramble or with shScramble, shICAM1#1 or shICAM1#3 following low-dose IFN-γ treatment. RT-PCR was performed using primers identified for the RBP-Jκ binding site in the *CD133* promoter region as follows: 5′- AGAGACTTCGGACTCGCTCT -3′, 5′- CACAGTGTTGGCCCATTTCC -3′.

### TCGA database

The TGCA NSCLC database was accessed via the UCSC Cancer Browser (https://genome-cancer.ucsc.edu). Gene expression based on TCGA RNA-Sequencing data is presented as the mean ± standard error of the mean (SEM) of triplicate determinants. Survival and correlation analyses of NSCLC patients were performed according to the TCGA RNA-Sequencing data and survival information. All analyses were performed using the R Package Survival.

### Statistical analysis

Data are shown as the mean ± s.d. and analyzed using Student’s t-test or χ^2^ test. A paired t-test was used for paired samples. Survival curves were plotted using the Kaplan-Meier method. Statistical analyses were performed using GraphPad Prism 7 software (GraphPad software, La Jolla, CA, USA). A value of *P*<0.05 was considered statistically significant.

## Author contributions

Y.Z., M.S. and Y.P. conceived the project. L.Y., F.L., B.Z., J.X. and D.Y. participated in the research design and coordination of the project. M.S. K.Z., S.C., J.Q., S.L. and T.S. performed the experiments. M.S. wrote the manuscript with input from Y.Z., Z.L., Y.P. and N.R.M. M.S. and K.Z. performed data analysis. All the authors read and approved the final manuscript.

## Acknowledgements

This study was funded by the National Key Research and Development Program of China (2016YFC1303501), the National Natural Science Foundation of China (Grant No. 81771781 and 81872410) and the National Key R&D Program (2018YFC1313400).

## The paper explained

### Problem

IFN-γ is conventionally recognized as an inflammatory cytokine that play a central role in antitumor immunity. Clinically, although has been used clinically to treat a variety of malignancies, low-level IFN-γ in the tumor microenvironment (TME) increases the risk of tumor metastasis during immunotherapy. Accumulating evidence has suggested that IFN-γ can induce cancer progression. The mechanisms underlying the controversial role of IFN-γ regulating tumor development remain unclear.

### Results

In this study, we firstly revealed a dose-dependent effect of IFN-γ in inducing tumor stemness to accelerate cancer progression in patients with a variety of cancer types. Mechanically, low-level IFN-γ endowed cancer stem-like properties via the ICAM1-PI3K-Akt-Notch1 axis, whereas high-level IFN-γ activated the JAK1-STAT1-caspase pathway to induce apoptosis in NSCLC. Inhibition of ICAM1 abrogated the stem-like properties of NSCLC cells induced by the low dose of IFN-γ both *in vitro* and *in vivo*.

### Impact

Our study first defines the role of low-level IFN-γ in conferring tumor stemness and clearly elucidate the distinct signaling pathways activated by IFN-γ in a dose-dependent manner, providing new insights into cancer treatment, particularly patients with low-level IFN-γ expression in the TME.

